# Constitutive, mosaic expression of TIE2 p.L914F during mouse development causes formation of venous malformation

**DOI:** 10.64898/2026.01.27.702061

**Authors:** Lindsay J Bischoff, Chhiring Sherpa, Sandra Schrenk, Elisa Boscolo

**Affiliations:** Division of Experimental Hematology and Cancer Biology, Cincinnati Children’s Hospital Medical Center, Cincinnati, OH, USA; Department of Cancer Biology, University of Cincinnati College of Medicine, Cincinnati, OH, USA; Department of Pediatrics, University of Cincinnati College of Medicine, Cincinnati, OH, USA

**Keywords:** TIE2 mutation, venous malformation, mosaicism, endothelial cells, blood vessels

## Abstract

**Background:** The hyperactivating p.L914F mutation in TIE2, a receptor tyrosine kinase that is essential for vascular development and function, has been found to drive sporadic venous malformation (VM). While germline or early developmental expression of the mutation is thought to be lethal, mosaic or somatic expression is expected to result in VM disease. However, this has never been shown experimentally. Therefore, we utilized a genetic murine model of TIE2 p.L914F to examine the effects of the mutation in the mosaic condition.

**Results:** Using an mTmG reporter mouse, we show that the *CMV-Cre* mouse line drives mosaic Cre recombination during early embryonic development. We then crossed *B6-Tg(Rosa26-TIE2*^*L914F*^)^*EBos*^ (*TIE2*^*L914F*^) mice to *CMV-Cre* mice, to drive mosaic expression of TIE2 p.L914F during development. The offspring of these mice did not have the expected Mendelian ratio of mutant to control animals, indicating that mutant mice experienced partial lethality during development. Furthermore, surviving CMV-Cre;TIE2^L914F^ mutant offspring developed a VM phenotype, with the formation of massively enlarged venous/capillary vessels in various tissues.

**Conclusions:** In this study, we show that mosaic embryonic expression of the VM-causative mutation TIE2 p.L914F causes partial embryonic lethality and the formation of a VM phenotype. This a novel *in vivo* model of mutant TIE2-driven VM disease and it illustrates that the extent of the mutational event during development will contribute to varying levels of severity in the VM phenotype.

## Introduction

Genetic disorders that appear congenitally or early during childhood can be caused by genetic mosaicism, in which a non-inherited mutation occurs during embryonic development, contributing to cell populations with different genotypes of the affected DNA loci or gene^1,2^. Mosaicism can therefore contribute to two phenotypic effects. First, the stage or cell type in which the mutation occurs determines the extent of the mosaicism. A mutational event occurring early during development will result in a higher percentage of cells carrying the mutation. The resulting phenotype may be extensive or localized depending on how many or which cell types are affected. Second, mosaicism may allow for mutations that would otherwise be embryonically lethal to persist and drive disease, as the mutational event may miss key developmental timepoints or effect too few cells to result in mortality^3,4^.

Mosaicism plays a role in vascular malformations, a group of disorders that result in the formation of malformed and dysfunctional blood and lymphatic vessels^4,5^. Vascular malformations are driven by genetic mutations that cause errors in normal vascular development, resulting in disease that appears congenitally or during childhood^6^. Venous malformation (VM), in which the diseased vessels consist of abnormally enlarged and dysfunctional veins, is the most common subtype of vascular malformation^7,8^. While VM appears in rare inherited cases^9^, most often it manifests as non-inherited, sporadic disease with single or multiple lesions^10^. The most common mutations that drive VM are located in the *TEK* gene, which encodes the TIE2 receptor tyrosine kinase^11,12^. TIE2 expression is enriched in endothelial cells (EC)^13^ and is important for normal vascular development^14,15^. More specifically, the TIE2 p.L914F mutation, which causes ligand-independent hyperactivation of the receptor^10,16,17^, occurs in up to 80% of TIE2-mutant VM. However, this mutation has only been identified in sporadic (somatic) cases of disease and never in inherited cases, indicating that it is likely lethal with germline transmission. Furthermore, the clinical presentation of TIE2 p.L914F-associated VM can be variable, with some patients experiencing localized lesions and some with more extensive impairment that can encompass entire limbs or other areas of the body^10,18,19^.

Based on this, embryonic TIE2 p.L914F mutation could persist when acquired in a mosaic fashion, which provides a rationale for the phenotypic variability in the extent of affected tissue or lesion size that is seen across patients. In this Research Letter, we report a novel murine model of VM based on embryonic mosaic expression of TIE2 p.L914F. This model recapitulates histological and phenotypical manifestations of patients’ VM.

## Results and Discussion

In our previous work, we established a novel murine line containing a Cre-inducible TIE2 p.L914F transgenic construct (*B6-Tg(Rosa26-TIE2*^*L914F*^)^*EBos*^, hereafter called *TIE2*^*L914F*^)^20^. Using these mice and the *Tg(Tek-cre)*^*1Ywa*^ (*Tie2-Cre*) line^21^, we showed that constitutive and endothelial cell (EC)-specific expression of mutant TIE2 p.L914F during early embryonic development caused embryonic lethality around E9.5^20^. However, this model does not allow for investigation of the phenotypic effects of the mutation in the mosaic or somatic condition.

Therefore, we crossed *TIE2*^*L914F*^ mice to the *B6*.*C-Tg(CMV-cre)*^*1Cgn/J*^ (*CMV-Cre*) line^22^ (**Figure 1A**), which was thought to induce constitutive Cre expression in all cell types early in development, but has recently been shown to have a variable and mosaic Cre recombination pattern^23,24^. Upon crossing *CMV-Cre* and *TIE2*^*L914F*^ mice we genotyped live offspring from the matings. Based on Mendelian inheritance patterns from the parental genotypes, the expected ratio of offspring genotypes (in the absence of any embryonic mortality) to be 50% control (*CMV-Cre*) offspring and 50% mutant (*CMV-Cre; TIE2*^*L914F*^) offspring. If the mutation was completely lethal, we would observe a complete lack of live mutant offspring (100% control, 0% mutant). However, our results showed an intermediate effect. Of the 106 total offspring, 90 (85%) were *CMV-Cre* controls and 16 (15%) were *CMV-Cre; TIE2*^*L914F*^ mutants (**Figure 1B**), revealing partial embryonic lethality in the TIE2-mutant group.

**Figure 1.**
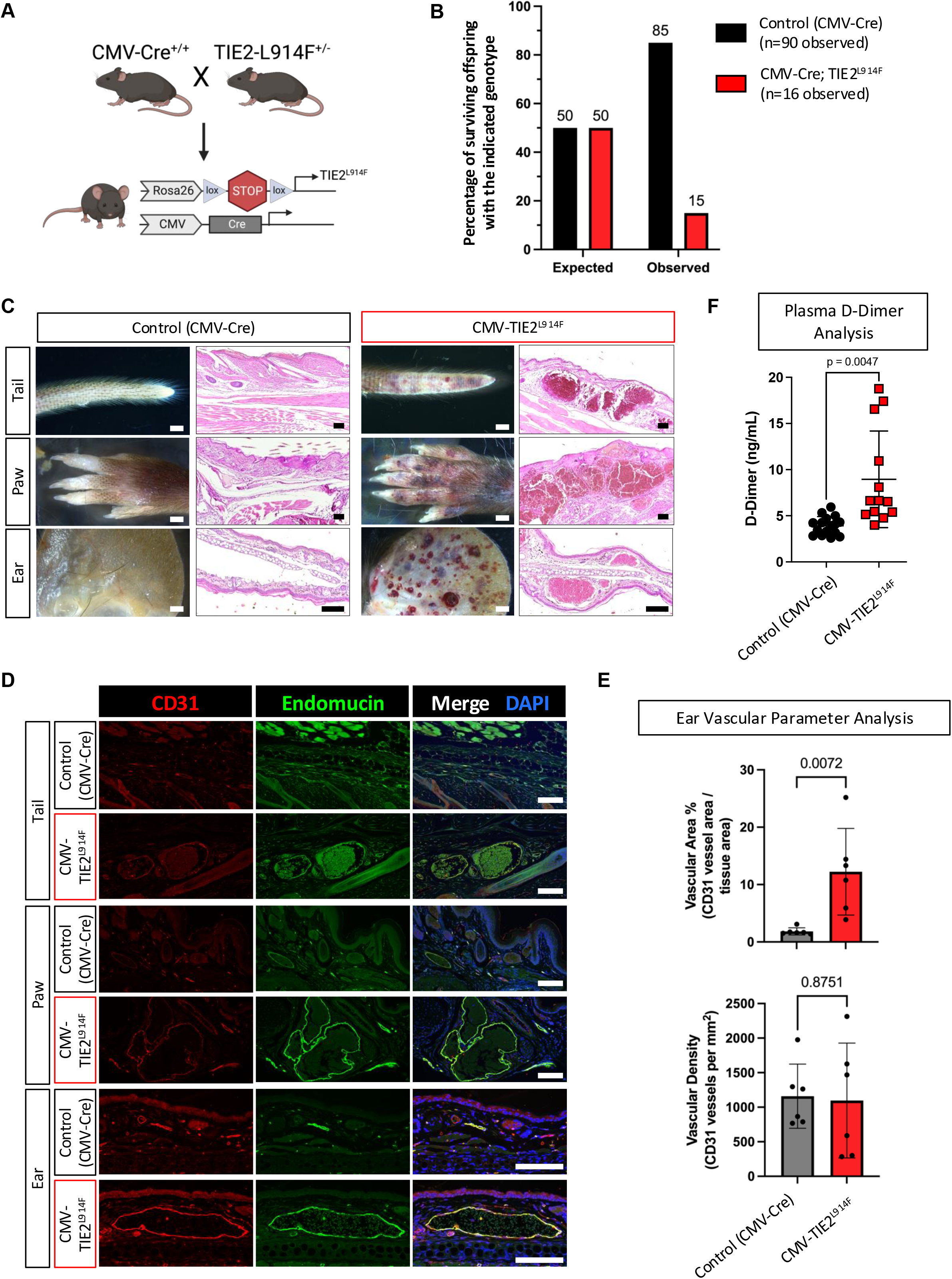
CMV-Cre driven mosaic expression of TIE2 p.L914F during development results in partial embryonic lethality and formation of ectactic blood vessels similar to VM. (A) Schematic of the CMV-Cre, TIE2^L914F^ breeding scheme and genotype of mutant *CMV-Cre, TIE2*^*L914F*^ offspring. Created with Biorender.com. (B) Expected and observed offspring genotypes from *CMV-Cre* x *TIE2*^*L914F*^ crosses. Expected genotype ratios were calculated from Mendelian inheritance patterns, assuming a parent with 2 alleles for *CMV-Cre* and a parent with 1 allele for TIE2^L914F^. Observed ratios were measured by genotyping 106 total offspring. Chi-square value for expected versus observed is 51.66 and p = <0.0001. (C, left columns) Photographs of gross tissues from control and mutant mice. Scale bar = 1mm. (C, right columns) Hematoxylin and eosin (H&E) staining of paraffin-embedded tissue sections. Scale bar = 100µm. (D) CD31 (red), Endomucin (green), and DAPI (nuclei, blue) immunostaining of paraffin-embedded tissue sections. Scale bar = 100 µm. (E) Quantification of vascular area and vascular density from CD31 and Endomucin immunostained ear sections. n = 6 mice per group. Mean±SD. Welch’s t-test. (F) D-dimer levels measured by ELISA in plasma of mice. Mean±SD. Welch’s t-test. p-values for all comparisons are indicated above the graph.

We further observed the *CMV-Cre; TIE2*^*L914F*^ mutant offspring that did survive. We found that, compared to control *CMV-Cre* siblings, the mutant mice developed visible and distinctive bluish /red vascular lesions, reminiscent of the VM patient phenotype. These lesions formed throughout the body, especially in the skin of the tails, paws, and ears (**Figure 1C, left columns**). Histological analysis of affected tissues revealed the presence of abnormal, massively enlarged blood vessels that disrupted normal tissue architecture, characteristic of VM lesional vessels (**Figure 1C, right columns)**. Immunofluorescence staining for CD31 (an EC-specific marker) showed the presence of massively enlarged CD31^+^ vessels that also were positive for Endomucin, a marker of venous/capillary vessel types (**Figure 1D**). We further quantified vascular parameters from ear tissue sections, revealing that vascular area is increased in *CMV-Cre; TIE2*^*L914F*^ mice, but vascular density is not (**Figure 1E**). Finally, VM patients, especially those with more extensive lesions^25^, can develop coagulopathy, one of the most severe and life-threatening complications of VM. Coagulopathy in VM is often detected by increases in plasma D-dimer levels^26^, which we also observed in *CMV-Cre; TIE2*^*L914F*^ mice (**Figure 1F)**. These data indicated that surviving TIE2-mutant offspring developed a vascular-specific phenotype, resulting in the formation of distinctive lesions consisting of massively enlarged and abnormal venous blood vessels, reminiscent of the patient VM phenotype.

Due to previous reports of mosaic *CMV-Cre* recombination^23,24^, we were intrigued by the possibility that the results we were observing were consistent with a model of genetic mosaicism. We therefore sought to verify the expression pattern of Cre-recombination in these mice under our experimental conditions. To do so, we crossed *CMV-Cre* mice to the *mTmG* reporter line^27^, in which Tomato fluorescent protein (mT) is expressed in all cells prior to recombination and green fluorescent protein (mG) is expressed after recombination, allowing for the visualization of both un-recombined and recombined cells (**Figure 2A**). As the *CMV-Cre* construct should be expressed early in embryonic development^22^, we collected E9.5 *CMV-Cre;mTmG* mouse embryos and performed whole-mount confocal imaging to determine patterns of recombination (**Figure 2B**). We found that recombination levels were highly variable in these embryos, with some exhibiting almost complete conversion to the recombined reporter (high levels of mGFP), some exhibiting very little recombination (high levels of mTomato), and some with mixed levels of both, indicating mosaic recombination. To quantify this variability, we measured the mean fluorescent intensity of both colors across the entire embryo and expressed the data as a ratio of mGFP to mTomato (**Figure 2C**), again showing that approximately one third of embryos had high levels of mGFP (high recombination), one third had intermediate levels of both mGFP and mTomato (mixed recombination), and one third had high levels of mTomato (little recombination).

**Figure 2.**
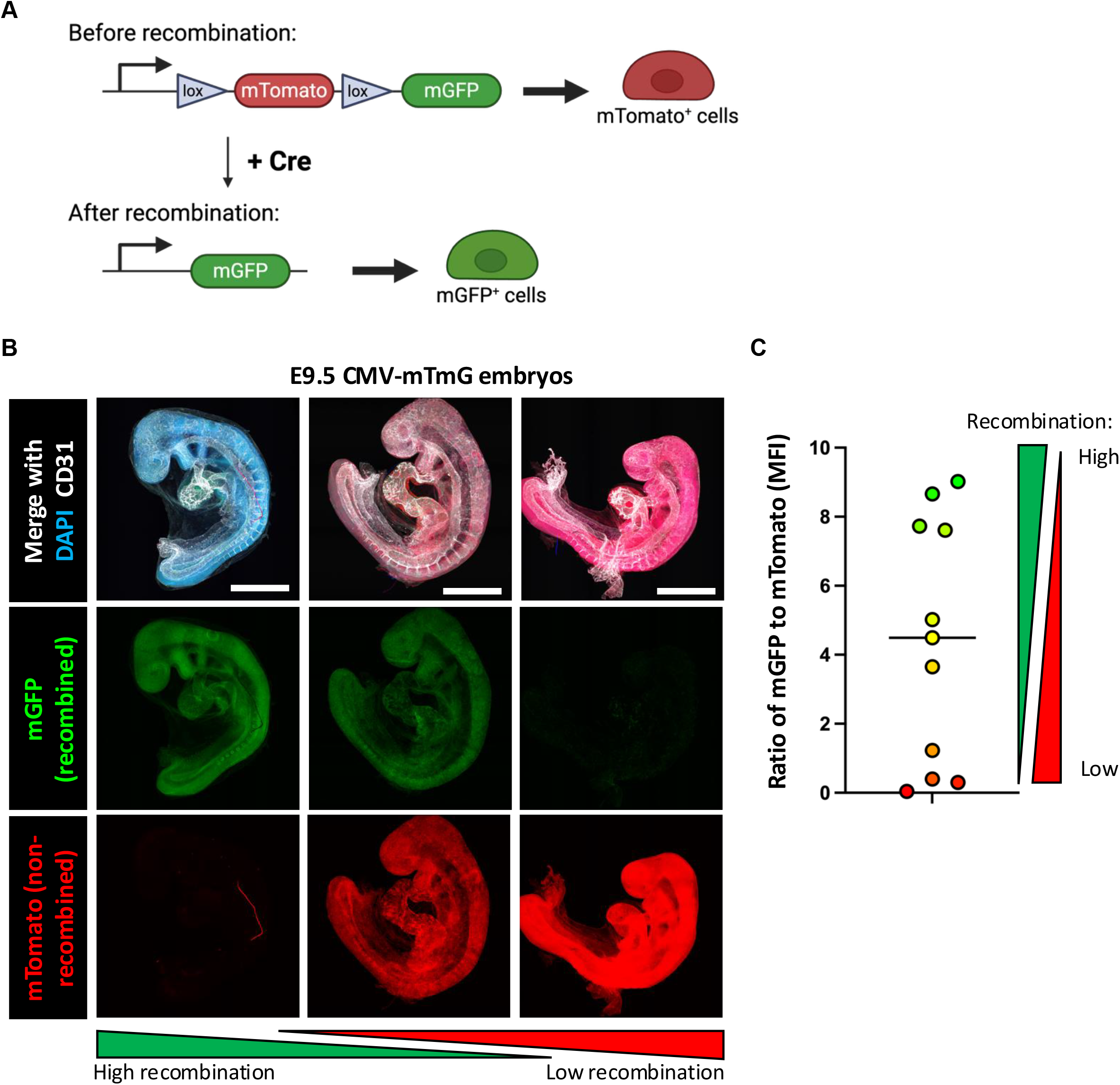
The CMV-Cre mouse line drives mosaic recombination during embryonic development. (A) Schematic showing the mT/mG reporter construct before and after Cre-mediated recombination and resulting fluorescence in cells. Created with Biorender.com. (B) Representative maximum intensity projections of tile-scan z-stack confocal images. Whole-mount E9.5 *CMV-Cre; mTmG* embryos were immunostained for the vasculature (CD31, white) and nuclei counterstained with DAPI (blue). Reporter fluorescence for mGFP is shown in green and fluorescence for mTomato is shown in red. Scale bar = 250µm. (C) For each embryo, the mean fluorescent intensity (MFI) of mGFP and mTomato was measured from maximum intensity projection images. Data is represented in the graph as the ratio of mGFP signal to mTomato signal. Each dot indicates one embryo. n = 11 embryos.

In total, these data indicate that *CMV-Cre;TIE2*^*L914F*^ mice that escape embryonic lethality eventually form vascular lesions reminiscent of VM. These results are supported by evidence that the *CMV-Cre* line promotes variable and mosaic transgene recombination^23,24^. Therefore, this study provides experimental evidence for the hypothesis that the timing and extent of transgene expression during development determines the phenotypic outcome of vascular malformations, with extensive recombination likely contributing to embryonic lethality and less extensive recombination (mimicking the mosaic condition) resulting in vascular-specific multifocal disease. We argue that this model most accurately mimics the random nature of genetic mutations in patients and recapitulates biological theory on the relationship of mosaicism and congenital disorders^1,4^.

## Experimental procedures

### Institutional permissions

All mouse experiments and procedures have been reviewed and approved by the CCHMC Institutional Animal Care and Use Committee (protocol number IACUC 2023-0025). All animals were cared for in accordance with guidelines from the National Institutes of Health. Studies were performed using both male and female mice.

### Mouse husbandry and experimental procedures

B6-Tg(Rosa26-TIE2^L914F^)^EBos^ (*TIE2*^*L914F*^) mice were generated, maintained, and genotyped as described previously^20^. Upon experiment endpoint or moribundity, mice were humanely euthanized according to guidelines established in our IACUC protocol by carbon dioxide inhalation and cervical dislocation.

To generate experimental offspring, male *B6-Tg(Rosa26-TIE2*^*L914F*^)^*EBos*^ mice were crossed with female *B6*.*C-Tg(CMV-cre)*^*1Cgn/J*^ (Jax stock No: 006054) (*CMV-Cre*) mice^22^ (to account for X-linked inheritance of the CMV-Cre allele). Surviving offspring were monitored for signs of lesion formation. *CMV-Cre* only littermate mice were used as controls for *CMV-Cre; TIE2*^*L914F*^ mutant mice.

To generate CMV-Cre reporter mice, male *Gt(ROSA)26Sor*^*tm4(ACTB-tdTomato,-EGFP)Luo*^*/J* mice^27^ were crossed with female *B6*.*C-Tg(CMV-cre)*^*1Cgn/J*^ (*CMV-Cre*) mice^22^. To obtain E9.5 embryos, mating pairs were combined and females were examined each morning for the presence of a vaginal plug, which was considered day E0.5. Pregnant females were euthanized before dissection of embryos, which were fixed in 10% neutral-buffered formalin prior to immunostaining and confocal imaging.

For D-dimer analysis, blood was collected from the inferior vena cava (IVC) by a 27-gauge syringe pre-loaded with 50uL of 0.105□M sodium citrate as anticoagulant. Samples were centrifuged at 1100□G for 10□min at room temperature. Plasma D-dimer levels were determined by ELISA (Asserachrom® D-Di, Diagnostica Stago, cat# NC9884012) following manufacturer’s instructions.

### Histology and immunostaining

Tissues were fixed in10% neutral-buffered formalin overnight at 4°C and then paraffin embedded and sectioned at 5µm for hematoxylin and eosin (H&E) staining or for immunofluorescence staining. For immunofluorescence, sections were deparaffinized with xylene and rehydrated through a descending ethanol series. Antigen retrieval was performed by boiling the slides in 0.01□M citric acid (pH 6.0). Immunostaining was performed with the following primary antibodies: CD31 (R&D #AF3628, 0.5µg/mL) and Endomucin (Santa Cruz #sc-65495, 4µg/mL). Counterstaining was performed with DAPI (4′,6-diamidino-2-phenylindole) (Invitrogen #D1306, 5□µg/mL).

### Confocal imaging and quantification

Z-stack images of tissue sections and whole-mount embryos were acquired on a Nikon A1 Inverted LUNV laser-scanning confocal microscope in the CCHMC Bio-Imaging and Analysis Facility (RRiD SCR_022628). Nikon NIS Elements software was used to create maximum intensity projections of the images.

## Tissue Section Vascular Parameters Quantification

Quantification of vascular parameters in ear sections was performed using Fiji (ImageJ)^28^ software. Vessels were identified by CD31 staining and manually outlined. Vessel density was calculated as the number of vessels in the total tissue area. Vascular area was calculated as the total area covered by vessels/vessel lumens over the total tissue area.

### Data collection and statistics

Prism 9.0 software (GraphPad) was used to generate graphs and perform statistical analysis. Data are presented as mean□±□standard deviation. Statistical significance between 2 groups was assessed by unpaired Welch’s *t*-test. Statistical significance is defined as *p* values less than 0.05. Schematics were created with Biorender.com.

## Acknowledgements and sources of funding

Research reported in this manuscript was supported by the National Heart, Lung, and Blood Institute, under Award Numbers R01HL117952 (E.B.), R01HL167700 (E.B.), and F31HL176101 (L.J.B.), part of the National Institutes of Health. The content is solely the responsibility of the authors and does not necessarily represent the official views of the National Institutes of Health. This manuscript is the result of funding in whole or in part by the National Institutes of Health (NIH). It is subject to the NIH Public Access Policy. Through acceptance of this federal funding, NIH has been given a right to make this manuscript publicly available in PubMed Central upon the Official Date of Publication, as defined by NIH. Additional funding supporting the study was provided by the American Heart Institute (AHA) Predoctoral Fellowship (https://doi.org/10.58275/AHA.24PRE1191403.pc.gr.190588) to L.J.B. We thank the Bio-imaging and Analysis Facility and the Comparative Medicine Division at Cincinnati Children’s Hospital Medical Center for providing state-of-the-art instrumentation, services, training, and education.

## Author contributions

L.J.B, S.S., and E.B. conceived the project. E.B. supervised the research. L.J.B. and S.S. performed the mouse experiments and sample collection. L.J.B. performed immunostaining and confocal imaging. L.J.B. and C.S. performed quantifications. L.J.B. wrote the manuscript and prepared the figures. All authors revised and commented on the manuscript.

